# Collapsed Site Coalescent Facilitates Detection of Adaptive Divergences Between Segmental Duplicates

**DOI:** 10.64898/2026.10.23.761411

**Authors:** Skylar LeMay, Arbel Harpak

**Affiliations:** Department of Computer Science, University of Texas at Austin, Austin, TX; Department of Integrative Biology, University of Texas at Austin, Austin, TX; Department of Quantitative and Systems Health Sciences, University of Texas at Austin, Austin, TX

## Abstract

Segmental duplications make up a substantial fraction of many genomes and are a recurrent source of evolutionary innovation, yet they have remained largely inaccessible to population genetic inference. One reason is that, until recently, their high sequence identity precluded reliable sequencing. Long-read sequencing now resolves population haplotypes across segmental duplications. Still lacking are appropriate population genetic methods to analyze population samples. A particular difficulty arises from frequent interlocus gene conversion (IGC), the copying of sequence from one duplicate into another. Such events correspond to coalescence of lineages across distinct sites—thereby violating a core assumption of standard coalescent theory and ancestral recombination graphs (ARGs). Here, we propose an approach that can facilitate ARG inference for segmental duplicates. We propose the Collapsed Site Coalescent (CSC), a model which treats each pair of paralogous sites as a single collapsed pseudo-site with a self-contained genealogy, so that the sequence of genealogies along a collapsed locus forms an ARG. Based on the CSC, we develop a haplotype-copying inference method that estimates IGC parameters from a population sample. In simulations, estimated parameters closely track true IGC rates. We further investigate the case where an adaptive difference between duplicates is sheltered by selection, protecting it from IGC that would otherwise erase the difference, a situation we call an *IGC haven*. When both IGC and selection are strong, our approach allows for the detection of IGC havens. In sum, this approach brings duplicated sequence within reach of population genetics inference, opening the way for tackling longstanding questions about the evolution of gene duplicates.

## 1 Introduction

“Segmental duplicates” are copies of a contiguous stretch of the genome, generated when errors in DNA replication, repair, or recombination duplicate a segment at the DNA level, inserting the copy either in an adjacent position to the original or at a distance. Segmental duplicates make up a large and rapidly evolving part of many genomes [1, 2]. In the first complete, telomere-to-telomere human genome, segmental duplications account for 7.0% of the sequence (218 Mbp), up from 5.4% in the previous reference assembly [1]. Long-read assemblies of 85 human genomes revealed a further 47 Mbp of duplicated sequence absent from the complete reference, as well as extensive variation in the structure and copy number of duplicates among individuals [3].

Segmental duplication is also the primary mechanism by which new genes arise [4], and although most gene duplicates are lost, those retained are a major substrate for evolutionary innovation [5, 6]. Examples include a duplicated pancreatic ribonuclease gene adapted to digest bacterial RNA in the foregut of a leaf-eating monkey [7]; a duplication of the Na^+^/K^+^-ATPase *α*-subunit gene that allowed neotropical frogs [8] to evolve resistance to dietary toxins while preserving pump function; and divergence between tandemly duplicated opsin genes underlying trichromatic color vision in Old World primates [9]. Which evolutionary paths duplicates follow, and what forces drive them, remain debated [4–6, 10].

Because of their high sequence identity, and the interlocus gene conversion (IGC) that helps maintain it, segmental duplicates have been largely inaccessible to population genetics. IGC is a form of recombination resulting in a non-reciprocal transfer of sequence from one duplicate to its paralog [11, 12]. Meiotic recombination is initiated by programmed double-strand breaks that are repaired using a homologous template, resulting in either a crossover or a gene conversion—a non-crossover in which a short tract of template sequence is copied onto the broken chromosome [12]. The template is usually the allelic sequence on the homologous chromosome, but when paralogous sequences are highly similar, as in segmental duplicates, the break can instead be repaired from the paralog, copying a tract from one duplicate into the other [12], which we call IGC. In humans, allelic gene conversion affects a given site with probability of about 6 × 10^−6^ per generation, with short tracts (tens to hundreds of base pairs) and a strong bias toward G/C alleles [13]. IGC between young duplicates is estimated to be an order of magnitude more frequent than point mutation, with a similar G/C bias and a rate that declines as paralogs diverge [14], and it accounts for at least 23% of the excess of single-nucleotide variants in segmental duplicates relative to unique sequence [15].

The parameters of IGC—including rate and tract length—therefore shape the evolution of a large fraction of the genome. By erasing differences between paralogs, IGC can cause duplicates to evolve in concert [16, 17], although how much it slows their divergence depends on how its rate changes as they diverge [14]. IGC contributes an estimated 7–8% of codon substitutions between teleost fish paralogs [18], and it introduces disease-causing alleles from pseudogenes into functional genes [12, 19]. How IGC rates vary along the genome and over time, and how they depend on sequence identity, distance between paralogs, and local recombination rate, remain largely open questions: experimental estimates of the rate have varied by orders of magnitude, likely reflecting such determinants, and most evolutionary estimates have been based on few loci [14].

IGC can also create a uniquely localized signal of natural selection. Because IGC continually erases differences between paralogs, surviving differences are likely maintained by selection—if a difference between duplicates is beneficial, conversion tracts that overwrite it will be purged. In such cases we would expect fewer IGC events and a lower IGC rate inferred (even when using a neutral model) at the immediate vicinity of the selected site, a signature we call an *IGC haven*. With high rates of IGC and strong selection, IGC havens can potentially pinpoint the targets of selection for divergence between duplicates at an unusually fine resolution. In the Na^+^/K^+^-ATPase genes of frogs that prey on toxic toads, IGC havens inferred from divergence between species identified both known resistance-conferring substitutions and previously unrecognized compensatory ones with pleiotropic effects [8]. As we discuss further in the **Discussion**, polymorphism data (within a single species) may offer an even richer resolution than divergence data on these signals, as it examines many generations cumulatively along the ARG, while being shaped by molecular processes and selective pressures reflective of recent evolution.

Studying the molecular evolution of segmental duplicates, understanding the drivers and consequences of IGC rate variation, and detecting IGC havens all require population genetic models that accommodate IGC. However, coalescent theory, and inference of the Ancestral Recombination Graph (ARG), rest on the assumption that the genome can be divided into sites, each with its own “marginal” tree-like genealogy [20–23]. ARG-based methods built on this assumption [22–26] have transformed the study of other regions of the genome [27–30], but IGC violates it: an allele in one duplicate can descend from the paralogous duplicate on the parent chromosome, so lineages at distinct sites in the genome can coalesce (**Fig. 1**).

**Figure 1.**
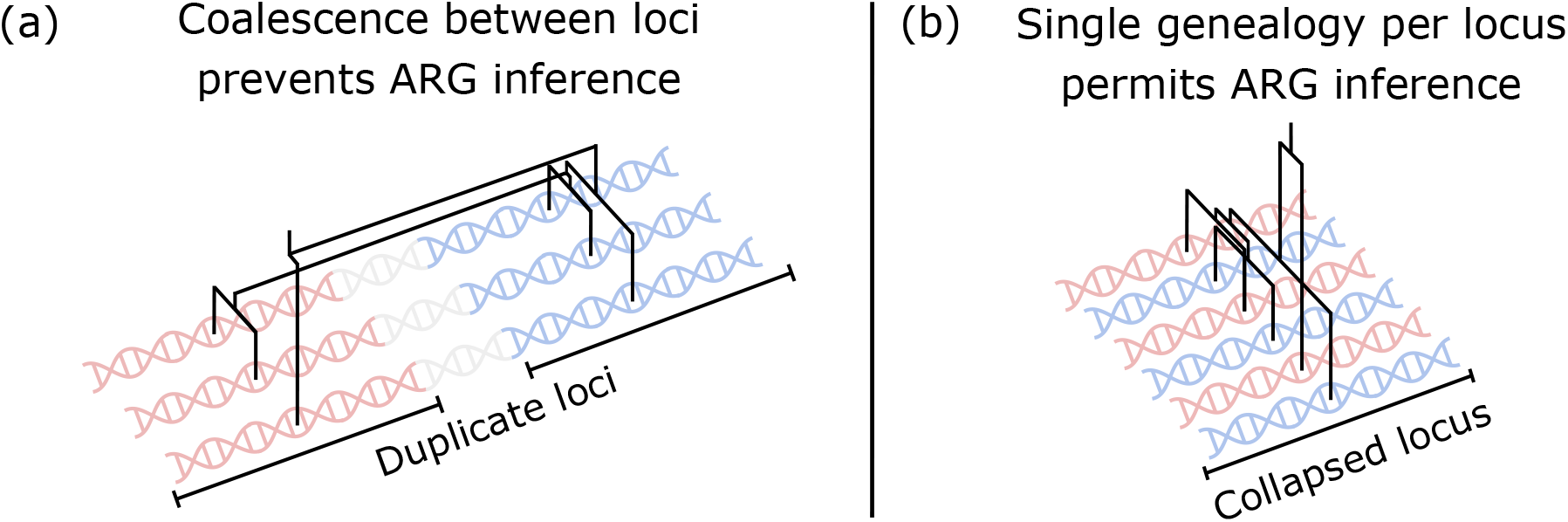
Collapsing duplicate loci facilitates ARG inference. **(a)** Genealogical relationships connecting three haplotypes at two paralogous sites within paralogous loci (shown in red and blue) representing segmental duplicates subject to the recombinational event interlocus gene conversion (IGC). IGC causes the coalescence of lineages across the two duplicates. **(b)** The same genealogical relationships as in **(a)** are represented as a single genealogy at a *collapsed site* within a *collapsed locus*. In some cases equal coalescence times are represented as slightly different in order to make the topology readable.

Here we follow in the footsteps of Innan [31] and Thornton [32], whose coalescent models of segmental duplicates have been used to derive expectations for summary statistics of polymorphism, and extend this approach to genealogical inference. We describe a coalescent model for a pair of segmental duplicates, the Collapsed Site Coalescent (CSC), which treats pairs of paralogous sites as a single *collapsed site* with its own genealogy (**Fig. 1**); the sequence of these genealogies along the *collapsed locus* forms an ARG for a pair of segmental duplicates. We then adapt the Li-Stephens (LS) copying model [33], a tractable approximation to the coalescent with recombination that has been highly successful in estimating crossover rates from polymorphism data [34–36], to the CSC. The resulting method, LS4Dups, estimates IGC parameters from the haplotype structure of a population sample, rather than from summary statistics of polymorphism [37] or from divergence between species [14]. In simulations, LS4Dups estimates track the per-site IGC rate closely, and the method detects IGC havens when both IGC and selection are strong.

## 2 Model

### 2.1 Collapsed site Coalescent

Population genetic inference methods based on the ARG [20, 38, 39] rely on the assumption that a chromosome’s allele at a particular site is copied (possibly with mutation) from the corresponding allele of a parent chromosome [20]. This means that all alleles at a site must share a common ancestor at the same site, which in turn means that a tree graph could accurately describe the genealogy at each site. The process of IGC violates this assumption, since the allele at a site within a duplicated region may sometimes be copied from the *paralogous* site of the parent chromosome. The effect of IGC on an ARG looks like lineages at one site branching onto, and coalescing with, lineages at the paralogous site (**Fig. 1**).

How can we adapt the standard coalescent and ARG models to describe this genealogical process? A first intuition might come from adopting existing coalescent models of population subdivision with migration [40]. One might model the two segmental duplicates as a sample from “red” and “blue” subpopulations of the same larger population at a single site, with IGC akin to migration between them. What such a model fails to capture is that segmental duplicates often result from tandem duplication on the same chromosome [16]. For a given site within the red locus and its paralogous site in the blue locus, an individual’s red and blue alleles at these sites will derive from the red and blue alleles of the same parent chromosome, unless a crossover occurs between the sites. This means that red and blue alleles can exhibit a type of correlated ancestry that has no equivalent in a migration model. We refer to red and blue alleles on the same chromosome as *coupled*. The possibility of crossover between the coupled alleles in a single lineage is also what prevents the representation of lineages in the form of a tree graph. To capture IGC and coupling within a coalescent framework, we developed the CSC model.

The core idea of the CSC is to treat paralogous sites as separate alleles from a single *collapsed site* (**Fig. 2a**). In this view, the paralogous sites collectively make up a *collapsed locus*. Since IGC only copies alleles within a collapsed site, there is no allelic exchange between distinct collapsed sites. This makes ARG inference possible by inferring self-contained genealogies at each collapsed site.

**Figure 2.**
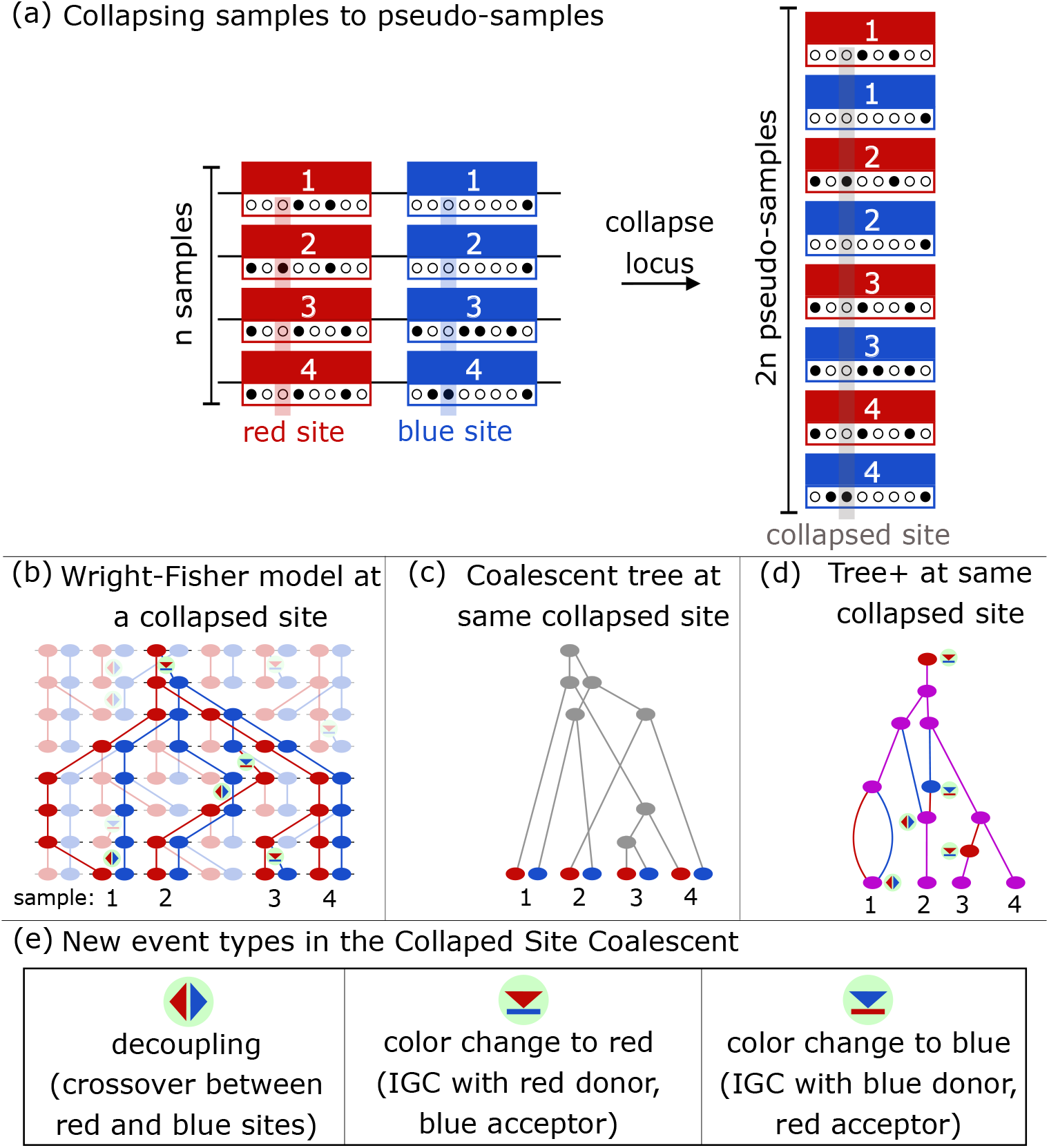
Collapsed Site Coalescent. **(a)** A sample of *n* chromosomes, each with two segmentally duplicated loci (left) may be *collapsed* and thought of as 2*n pseudo-samples* from a single collapsed locus. In this view, rather than IGC representing allele exchange between distinct sites (red and blue), IGC represents a form of coalescence within a single collapsed site. One particular collapsed site is highlighed in grey on the right, with the two sites it corresponds to highlighted in red and blue on the left. **(b)** An example of the Wright-Fisher model at a single collapsed site. Time flows downward from past to present. The highlighted nodes are ancestral to the 4-chromosome sample taken in the present, and correspond to the Collapsed Site Coalescent (CSC) on this sample. Coloration throughout represents which site ancestral material is found in—the red site, the blue site, or both (purple). **(c)** The coalescent tree between the sampled chromosomes (the nodes at the bottom of **(b)**). Branches are colored grey to indicate that this structure captures only the relatedness between leaves and not ancestral site. This structure is implied by the tree structure shown in **(d)** but contains strictly less information, as it does not capture the coupling of lineages (purple branches in **(d)**). **(d)** The tree+ corresponding to **(b).** The coloration of branches indicates location of ancestral material, with purple branches representing red and blue loci in the same ancestral chromosomes, both carrying ancestral material. **(e)** The new event types present added the CSC (on top of coalescence which is already present in the standard coalescent).

We begin by defining the CSC for a red and blue locus resulting from a segmental duplication^1^. In practice, many segmental duplicate (SD) families include more than two members, but we restrict our attention here to size-2 families for simplicity. We discuss potential extensions to larger gene families in the **Discussion**. A site in the red locus has a paralogous site in the blue locus which may exchange alleles through IGC. We will consider these sites as a single collapsed site (**Fig. 2a**), and define a coalescent model based on an adapted Wright-Fisher process for a collapsed site.

In the Wright-Fisher process for a single site, a diploid population is modeled as a set of 2*N* nodes for a parameter *N* representing population size. Each generation is generated by sampling the previous generation’s 2*N* nodes uniformly with replacement. Each of the 2*N* alleles have a fixed probability of mutation to a new allele (infinite sites mutational model; [41]). This model’s predictions converge with that of the standard discrete-time coalescent model, wherein we consider *n* tracked lineages in a population of *N* diploids, beginning with our sample at time zero. In each generation, each pair of samples derives from a common ancestor (“coalesces”) in the previous generation with probability 1 / 2*N*, and a relatedness tree is reconstructed by proceeding backwards, merging samples when they coalesce, until a most recent common ancestor for all samples (“root” of the coalescent tree) is found.

A collapsed site is modeled by a Wright-Fisher model for a single collapsed site, or two paralogous sites *s*_*R*_ and *s*_*B*_ simultaneously. In this model, each chromosome holds a pair of paralogous alleles. Due to linkage between *s*_*R*_ and *s*_*B*_, a chromosome will copy both its red and blue alleles from the same chromosome in the previous generation, except in the case of a crossover between *s*_*R*_ and *s*_*B*_, which occurs with fixed probability. Due to IGC, a red allele will copy the blue allele from the parent rather than the red with some fixed probability, and a blue allele will copy the red allele with the same probability (**Fig. 2b**).

The CSC (**Fig. 2b**) is a backward-in-time version of this collapsed site Wright-Fisher model. Implicitly, the CSC tracks linkage and crossover between the two uncollapsed sites, and also incorporates IGC events. It begins with a set of terminal nodes, each holding red and blue alleles which are coupled. The coupling notion in the CSC means we lose exchangibility across all lineages—different lineages carry different types of ancestral alleles. However, we introduce the addition of a color label to each lineage to maintain exchangibility among lineages of the same color.

Namely a purple lineage corresponds to a chromosome that has both of its paralogous alleles ancestral to the leaves, and we say it contains a red and a blue *sub-lineage*. A red (blue) lineage consists of only a single red (blue) sub-lineage. Each present-day sampled chromosome corresponds to a terminal node which is the start of a purple lineage (purple terminal nodes in **Fig. 2d**). A *decoupling* occurs when, due to a crossover between the paralogous sites, a chromosome in a purple lineage copies its two alleles from two different parent chromosomes, splitting the red and blue sub-lineages associated with the purple lineage into separate lineages. The color labeling makes IGC events affecting the collapsed site correspond to a *color change* for the lineage they occur in. The color change will switch a lineage’s color to the color of the donor. For example, a color change to blue occurs when underlying IGC uses a blue donor at the expense of a red recipient (**Fig. 2d**). A color change on a purple branch means that the sub-lineages associated with the lineage coalesce into one sub-lineage. A *coalescence* (of lineages) occurs when two chromosomes in one generation descend from the same parent chromosome in the previous generation. When two lineages coalesce, associated sub-lineages sharing the same color will coalesce. If a red lineage and blue lineage coalesce, a *coupling* occurs, in which case the parent will be a purple lineage. The process ends when only one sub-lineage (either red or blue) remains. As in standard coalescent theory (and unlike the multiple merger coalescent [42]), we assume the probabilities of multiple events—simultaneous color changes, couplings, decouplings or coalescences in a single generation, or simultaneous coalescences of more than two lineages—are negligibly small.

An instantiation of the CSC generates a directed acyclic graph whose leaves correspond to the present samples, whose internal nodes correspond to coalescences, decouplings, or color changes, and whose colored branches correspond to colored lineages in the CSC. We refer to this graph structure as a *tree*+ (**Fig. 2d**), to emphasize that it is the equivalent of a marginal tree at a single site in the standard coalescent, but that it does not itself have a tree structure if it includes decoupling nodes, as these have two parents (**Table 1**). In analogy with sub-lineages, we consider purple branches to contain red and blue *sub-branches*.

**Table 1.** Events in the Collapsed Site Coalescent. *k* indicates the total number of active lineages, and *k*_*R*_, *k*_*B*_, *k*_*P*_ the total red, blue, and purple active lineages respectively. All rates indicate the rate parameter of the exponential waiting time distribution corresponding to the event in question, and are given in coalescent units, of reciprocal of 2*N* generations. Because we assume that the probability of multiple events in a single generation is negligible, the per-generation event probabilities are directly proportional to the rates shown.

| event type | lineages below / above | state change | rate |
| --- | --- | --- | --- |
| leaf | 0 / 1 | new purple | 1 |
| decoupling | 1 / 2 | purple → red, blue | $k_P R$ |
| color change (red donor) | 1 / 1 | purple → red<br>blue → red | $(k_R + k_P)C$ |
| color change (blue donor) | 1 / 1 | purple → blue<br>red → blue | $(k_B + k_P)C$ |
| coalescence | 2 / 1 | purple, purple → purple<br>purple, red → purple<br>purple, blue → purple<br>red, red → red<br>blue, blue → blue | $\binom{k_P}{2}$<br>$k_P k_R$<br>$k_P k_B$<br>$\binom{k_R}{2}$<br>$\binom{k_B}{2}$ |
| (coupling) | | red, blue → purple | $k_R k_B$ |

A tree+ embeds an underlying tree in which every sub-branch is represented as its own branch, and color changes on purple branches are coalescences of these sub-branches (**Fig. 2c**). This representation does not contain information about ancestral coupling and so is strictly less informative than the tree+.

The lengths of the branches in the tree are distributed according to the CSC rates (**Table 1**). We will use time units of 2*N* generations where *N* is the population size (the standard “coalescent units” [21]) and, as in the standard coalescent [21], we will assume the population size is large enough that we may use a continuous-time approximation, with the wait times to events modeled with continous exponential distributions. Purple lineages decouple at rate *R*, and if there are *k*_*P*_ such lineages, then the decoupling rate for *k*_*P*_ purple lineages is *k*_*P*_ *R*. Color changes to red (blue) occur at rate *C* per non-red (non-blue) lineage, since purple lineages contain one blue (red) lineage offering an opportunity for such a color change. Therefore, for *k* lineages of which *k*_*P*_ are purple, the total color change rate is 2*k*_*P*_ *C* + (*k* − *k*_*P*_) *C =* (*k* + *k*_*P*_) *C*. Any pair of lineages (regardless of color) experiences coalescence at rate 1 (due to the choice of length units). For *k* lineages, coalescence occurs at rate 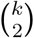. Based on these per-generation probabilities, we may derive the expected waiting times between events, which determine the time to most recent common ancestor (TMRCA) and the expected total sub-branch length (TSBL). We introduce TSBL as an analog for the total branch length within a standard coalescent tree [43], in order to count the red and blue sub-branch components of purple branches separately. For instance, the rate of mutation across the full tree will be proportional to TSBL, rather than total branch length, since both red and blue sub-branches may experience mutations. The expected values may be computed exactly by a dynamic programming algorithm applied to the state space of possible configurations of active lineages (i.e. possible combinations of color counts *k*_*P*_, *k*_*R*_, *k*_*B*_). One possible approximation comes from thinking of the CSC as being composed of two separate trees, a red and a blue, which must each coalesce to the root before the two coalesce together, and which have a net zero expected flow of lineages between them assuming symmetrical IGC rates. In this approximation the dynamic programming algorithm is only needed for the waiting time from the semifinal to the final coalescence. If *E* [*TMRCA* ∣ *n, R, C*] and *E* [*TSBL* ∣ *n, R, C*] are the expected TMRCA and TSBL for a sample of *n* haplotypes with *R* and *C* the per-site decoupling and color change rates, then the approximation is

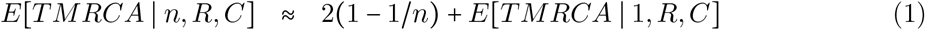

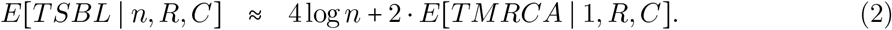

This approximation neglects the effects of color changes on purple lineages occurring below the most recent common ancestor, which reduce the total number of sub-branches. When color changes are very rare relative to coalescence (*C* << 1), this simplification will have a small effect, since color change will be less frequent lower in the tree+, but when color change is common relative to coalescence, this will cause the approximation to be biased towards overestimation, particularly for TSBL due to the reduction in sub-branch count (**Fig. 3**). We discuss details in **Supplementary Note Section S3**. We used the expected TMRCA in our tests described in **Validation** to determine the number of generations to run the forward simulation.

**Figure 3.**
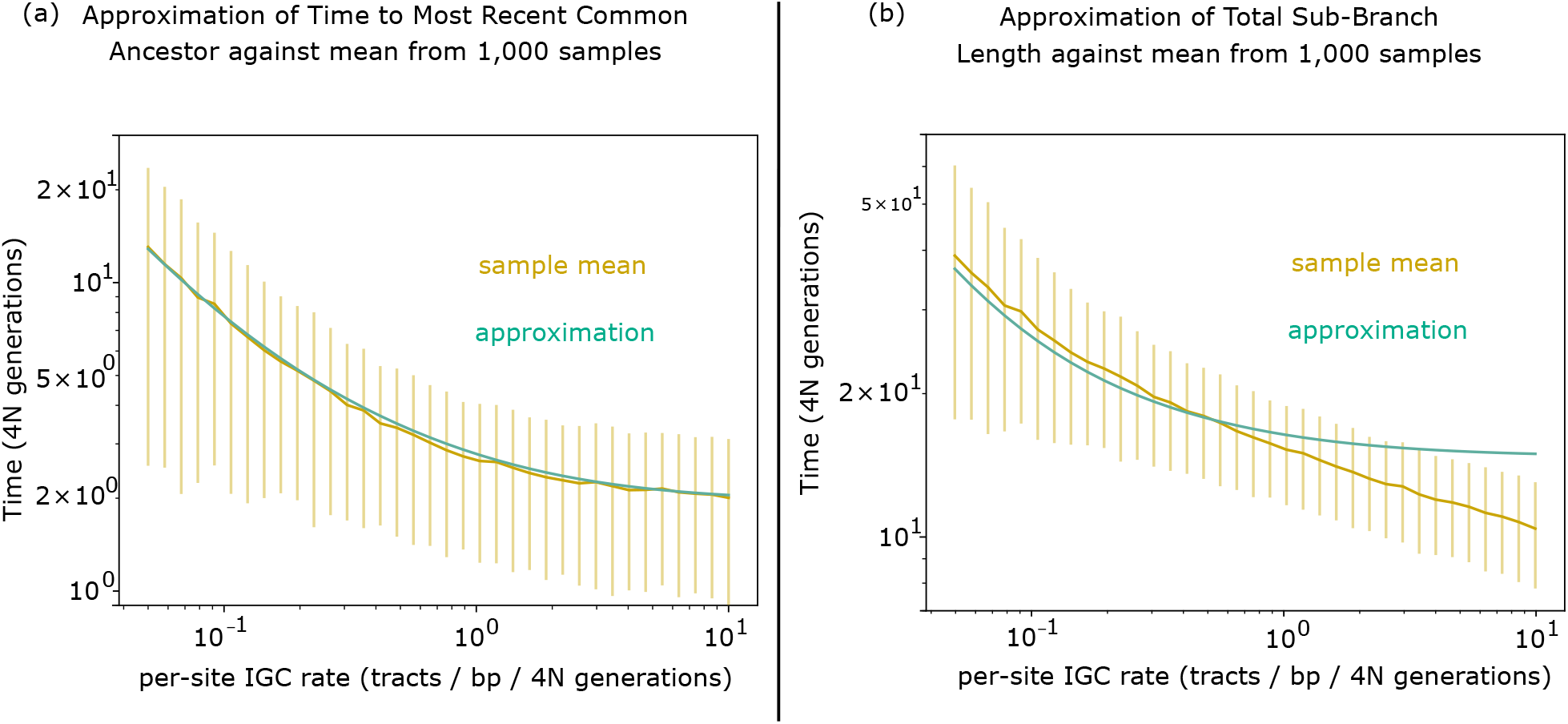
Approximations for total time to root and branch length in the CSC. Comparison of the approximations in **Eq. 1** and **2** for the expected values of two statistics of a tree+ —the Time to Most Recent Common Ancestor (TMRCA; total time elapsed from the most recent common ancestor of) (**(a)**) and expected Total Sub-Branch Length (TSBL) (**(b)**)—against their true distribution across 1,000 simulated tree+ s generated according to the CSC with varying per-site IGC rate *C* (CSC color change rate parameter). The per-site decoupling rate *R* was set to 1. We chose this value since the per-site decoupling rate is *R* = 4*Nr*Δ where *N* is the population size, *r* is the average per-generation recombination rate between the paralogs, and Δ is the distance between paralogs, and in our later inference simulations(**Fig. 5** and **Fig. 6**), we chose *r* ~ 10^−8^, *N* ~ 10^2^, and Δ ~ 10^6^. The sample means are shown in a gold line with error bars showing ±1 sample standard deviation for each sample (they appear off-center due to the log-scaling but are centered), with the approximation shown by a teal line. For TMRCA (**(a)**), the approximation was very close to the sample means. For TSBL (**(b)**), the approximation was biased upward for large *C* as discussed in the main text.

### 2.2 LS model for non-duplicated sequences

We wish to use population genetic inference methods on polymorphism data from a collapsed locus. As a starting point, we look to best-practice genealogical inference approaches for non-collapsed sites. Methods that tried to infer ARGs by integrating over all possible trees [22] have proven intractable computationally. For this reason, heuristics have been developed for approximating the ARG, for example by approximating the likelihood of haplotype data given population-genetic model parameters [22–26, 44]. The LS copying model [33] is a component of several such ARG inference methods [23–25]. The idea of the LS model is to iteratively generate haplotypes as mosaics of other haplotypes (“templates”) with occassional modifications. The rate of modifications is linked to the mutation rate. The rate of changes in template is linked to the crossover rate.

The LS model broke new ground and proved very useful in inference about molecular parameters such as crossover rates [45, 46]. By “molecular” crossover rate, we mean the expected number of crossovers per base pair per meiosis in a genomic region. One key advantage of this model is that the copying process closely tracks aspects of the coalescent process, whose model parameters, in turn, closely adhere to molecular parameters. Namely, the bootstrapping of new focal haplotypes from existing ones is akin to the branching in of a focal sample into marginal trees of previous samples. When a new focal sample is added to the coalescent tree for a site *j*, its first coalescence with the rest of the tree will add an ancestral node to some branch that includes the template sample as a descendant. The *j*’th allele of the focal sample will be the same as the template’s *j*’th allele, unless a mutation occurs along the path from the ancestral node to the focal sample or the template. At the adjacent site *j* + 1, if no crossover between sites *j* and *j* + 1 changes the tree topology, the first coalescence node will be the same. If a crossover occured, a new coalescence node is sampled. Nevertheless, the LS model does not build explicit marginal coalescent trees nor does it estimate marginal trees’ branch lengths, and so methods incorporating it usually include additional steps towards full ARG inference [23–25].

The LS model generates a matrix of haplotypes *h*_1_, …, *h*_*n*_. Each haplotype holds ℓ alleles, representing the allelic states at the ℓ single nucleotide polymorphisms (SNPs) within a DNA segment of total length *L* base pairs. These SNPs are modeled as biallelic, so that *de novo* mutations at the same site result in the same derived allele (encoded as 1), and back mutations restore the ancestral allele (encoded as 0). Every individual haplotype is generated by a hidden Markov model (HMM) with the identity of the template as the hidden state and the allele as the observed state. **Algorithm 1** is the LS generative algorithm. This model can be used to generate haplotype data, but can also be used to evaluate an approximate likelihood of a particular haplotype matrix under particular parameters, using the forward-backward algorithm [47]. The estimated likelihood may vary with respect to permutations of the haplotypes, as it depends on the randomly sampled order *k*_2_, *k*_3_, …, *k*_*n*_ and randomness in the initialization of *h*_*k*1_. In the original work of Li and Stephens, the likelihood is averaged over a small number of random permutations (20 permutations), and it is validated empirically that the number of random permutations does not strongly affect model behavior [33]. The LS parameters reflecting effective crossover rate (*ρ*_*j*_/*k* in the *j*’th region) and mutation rate 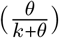 decrease as the number of templates *k* increases. This is what is predicted by the coalescent model—more tree branches means a shorter expected time to coalescence, which reduces the amount of crossover and mutation expected between the focus or template and the common ancestor of the two—but the expressions are heuristic: the inversely proportional scaling of crossover probability with respect to *k* is not derived directly from the coalescent. The value of the population mutational parameter *θ* is chosen so that the expectation of the number of mutations that occurred in the tree at each polymorphic site is *θ* · *L* = 1, a choice based on the infinite sites assumption but which does allow for multiple mutations at a single site.

#### Algorithm 1

LS generative model

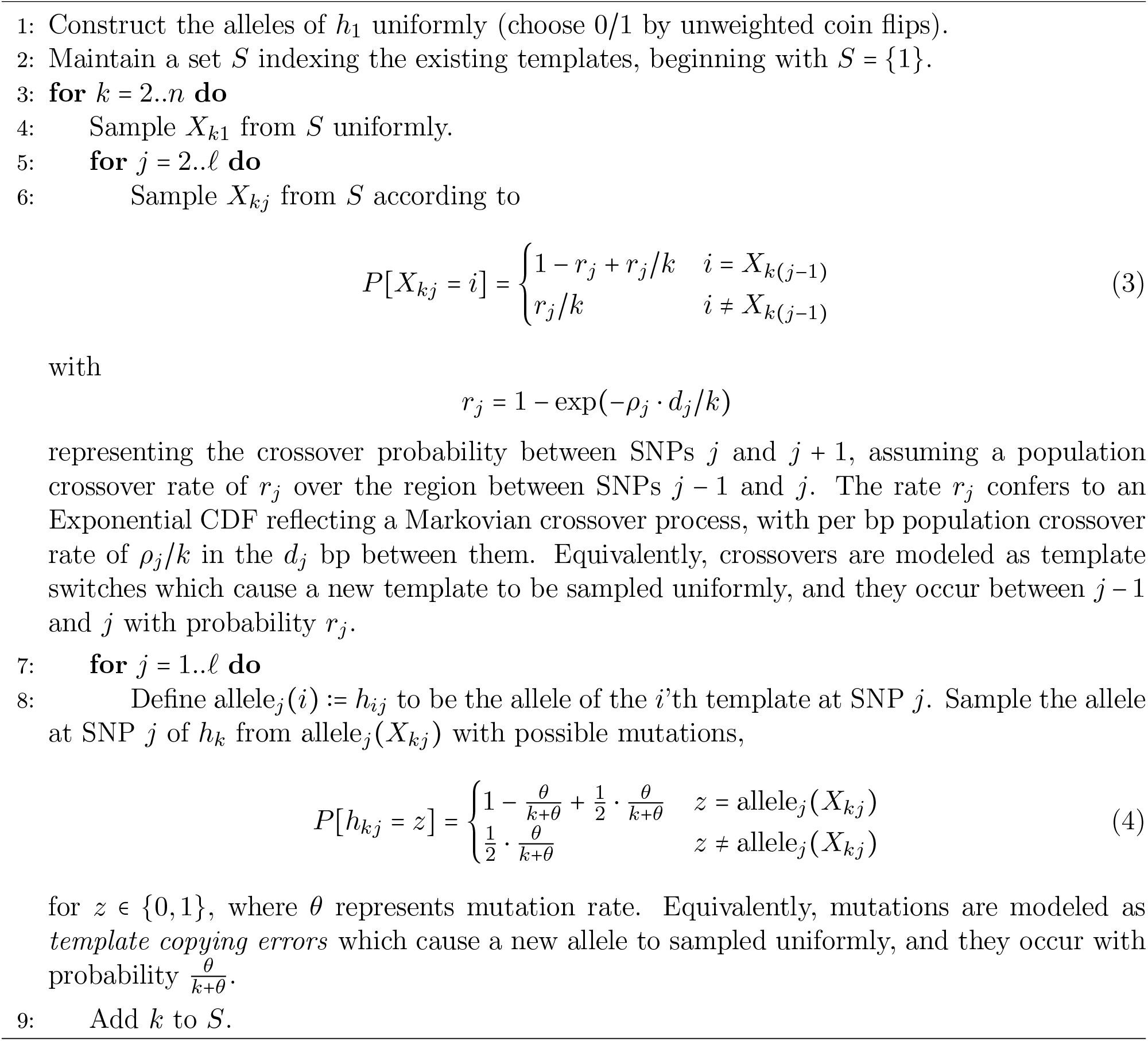

### 2.3 Modified LS model for segmental duplicates

We propose a new algorithm, LS4Dups, that adapts the LS model to the CSC (**Fig. 4**). This model is motivated by the same notion of iteratively adding new samples to a genealogy, but the corresponding iteratively updated genealogy now has the form of a tree. As before haplotypes are generated from previously generated haplotypes, but each sample now contains two pseudo-samples, each with its own haplotype. Each individual haplotype 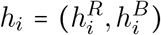 (we represent the red and blue components of the *i*’th haplotype *h*_*i*_ by 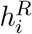 and 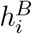 respectively) is generated independently from existing haplotypes, and the two pseudo-samples in a sample are generated in a random order. It is possible for a haplotype to copy from another haplotype of a different color, which in the underlying tree+ corresponds to a color change occurring on the lineage from the focal pseudo-sample to its common ancestor with the template.

**Figure 4.**
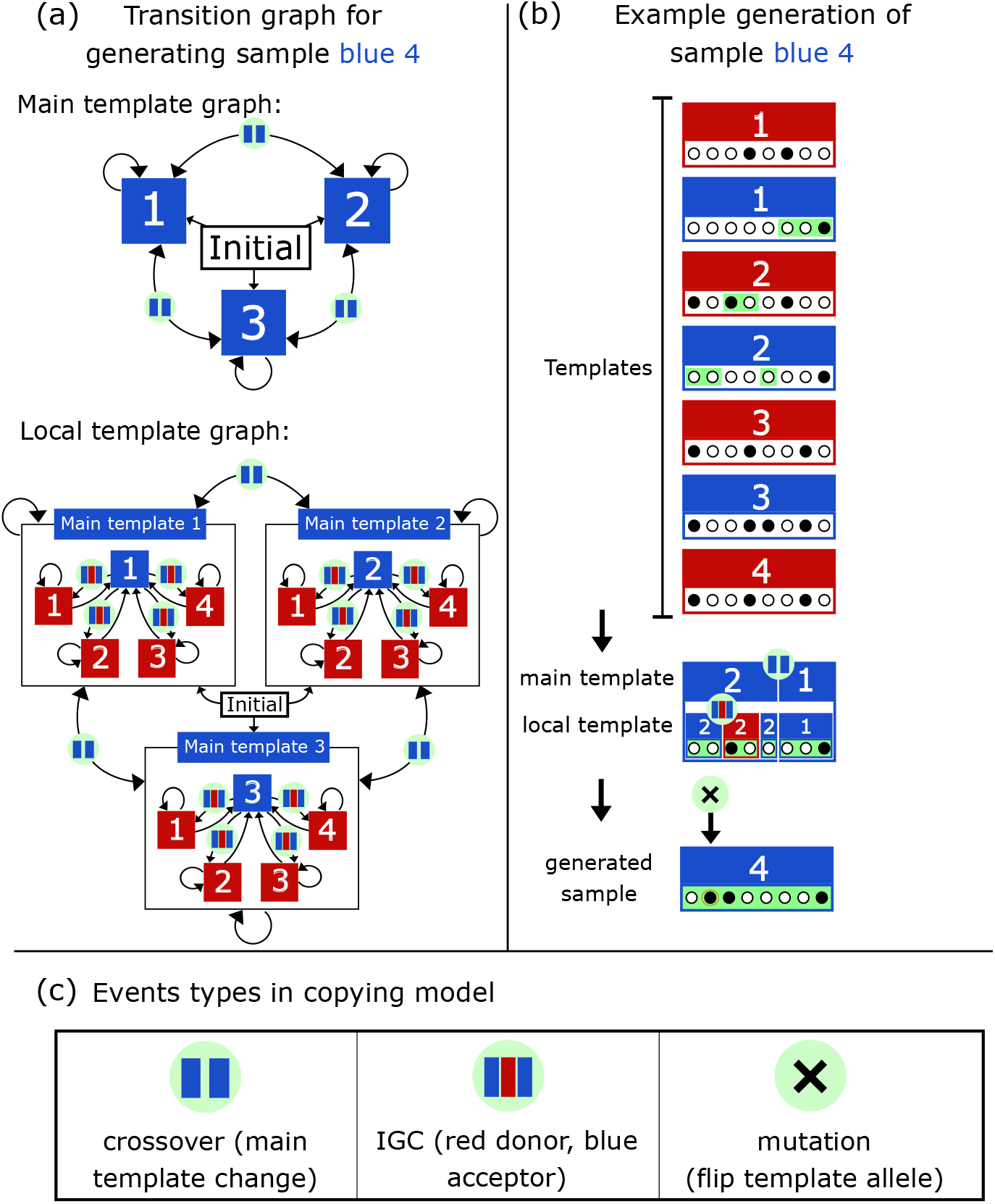
LS4Dups copying model. We estimate IGC parameters using a copying model HMM adapted from [33]. As in the LS HMM, sample *i* is generated from the previous samples from 1 to (*i* − 1). Here we show an example of generating the blue locus of the 4th sample (blue 4), from blue and red 1-3, as well as its own paralog red 4. **(a)** The transition graph between hidden states in our copying model for generating blue 4. To generate a sequence, first a sequence of main templates is generated via the *main template graph* which includes only templates of the same color as the focal pseudo-sample (here blue). Then within each contiguous subsequence of fixed main template, a local template sequence is generated, which may include tracts of other colors (here red). The even arrow thickness does not represent relative probability; for instance, transitions to red 4 will be more likely than other IGC transitions because of the preferential weight given to blue 4’s paralog in IGC tracts. **(b)** An example of sequence generation via the method shown in **(a)**. The main template sequence involves a single template switch (crossover between paralogous sites) from blue 2 to blue 1, but in generating the local template sequence, a tract from red 2 is inserted within the main template blue 2 region. After this sequence is generated, it is copied with a probability of mutation at each position. As in the original LS model we model polymorphisms as biallelic, so that a mutation is simply a flip of the allelic state of the template allele. **(c)** The event types involved in the copying model, and their effects on the copying process.

We include a parameter *α*_*k*_ (*k* the number of samples) that specifies a higher probability for a haplotype to copy from its own paralog haplotype (if it has previously been generated) than from the other different-color haplotypes. From the perspective of adding a focal pseudo-sample *f* into a tree+ on *k* existing samples which coalesces at a point *a*, the parameter *α*_*k*_ captures the possibility of color change on the branch from *f* to *a*, while the *f* lineage is in its original coupled state—there has no decoupling between the leaf and the IGC. Its value is determined by the relative sizes of color change rate and decoupling rate. The paralog will coalesce with the focus first if that lineage experiences a conversion before a decouping or a coalescence. The expected time until the focus’s first coalescence is the same as in the standard coalescent, namely 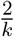 where *k* is the number of samples. A conversion will occur along this branch with probability 1 − exp −2*C* 2 / *k*) = 1 −exp (−4*C / k*). Such a conversion occurs before a decouping with probability 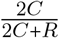. Therefore we set

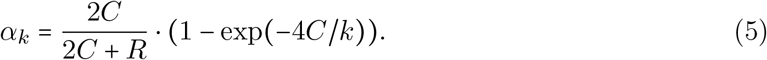

In our implementation we set the mutation parameter to 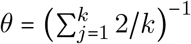 as in the standard LS model.^2^ **Algorithm 2** shows our adaptation of the LS model.

Although our model LS4Dups does involve crossover between adjacent sites in the same way as the standard LS model, the only parameter we include that would correspond to the decoupling parameter *R* from the CSC is the paralog preference *α*_*k*_, which does not capture the types of correlation that may exist outside of color changes (e.g. with very weak decoupling, if 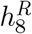 copies from 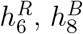 is more likely to copy from 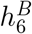). To fully include decoupling in this way, we would need to model the correlation between the template choices of 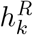 and 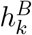, with weaker decoupling leading to more correlated template choices. We do not include this in our model for reasons of computational complexity. Namely, jointly modeling the template choices of 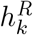 and 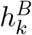 would require a larger state space in the hidden Markov model, of order *O*(*n*^2^) for states representing pairs of templates rather than *O*(*n*) for states representing single templates, which would impose significantly larger computational demands. We note that the correlation of red-red with blue-blue coalescences contains information about the average crossover rate between paralogous sites which our model does not incorporate.

#### Algorithm 2

LS4Dups generative model

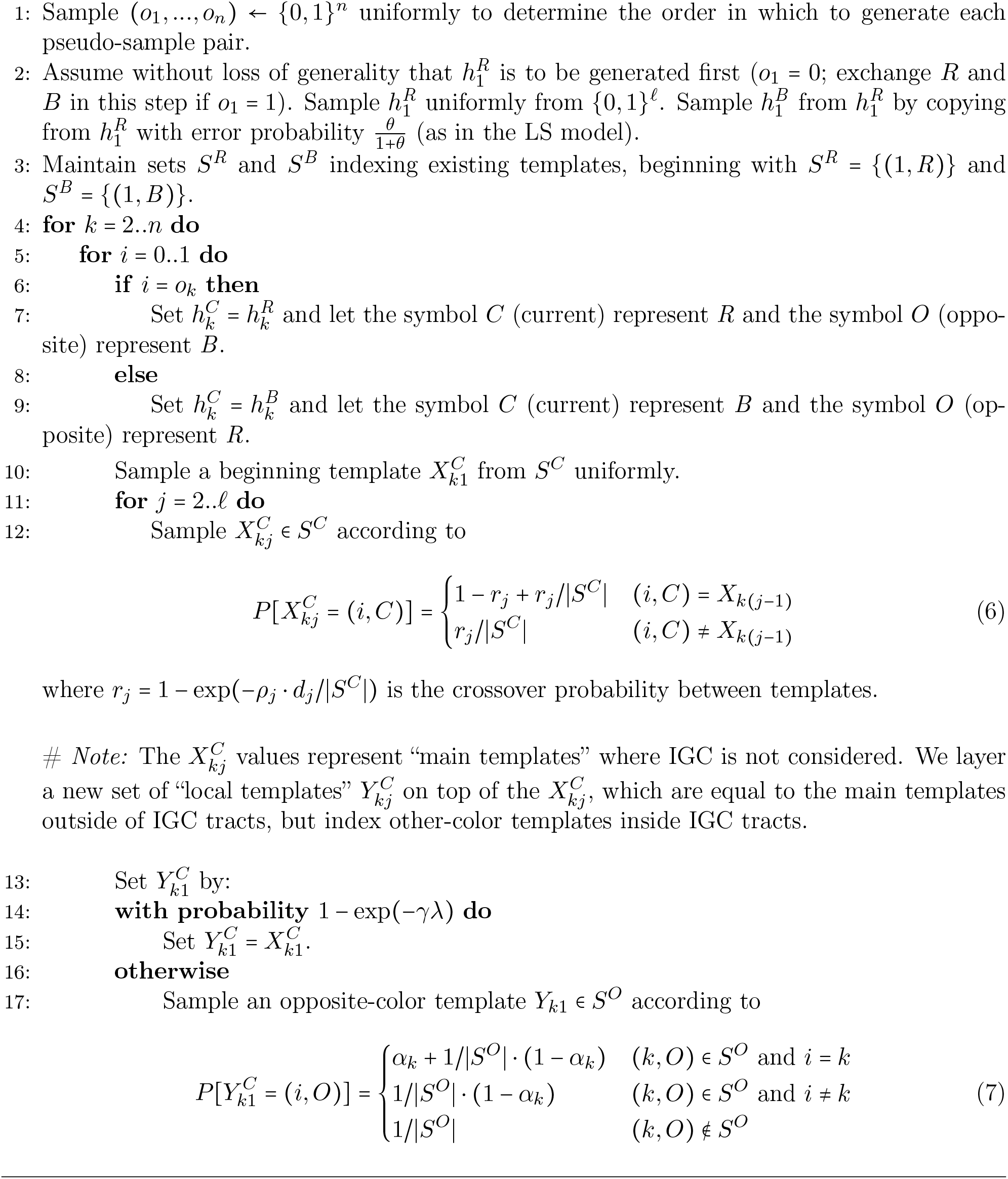
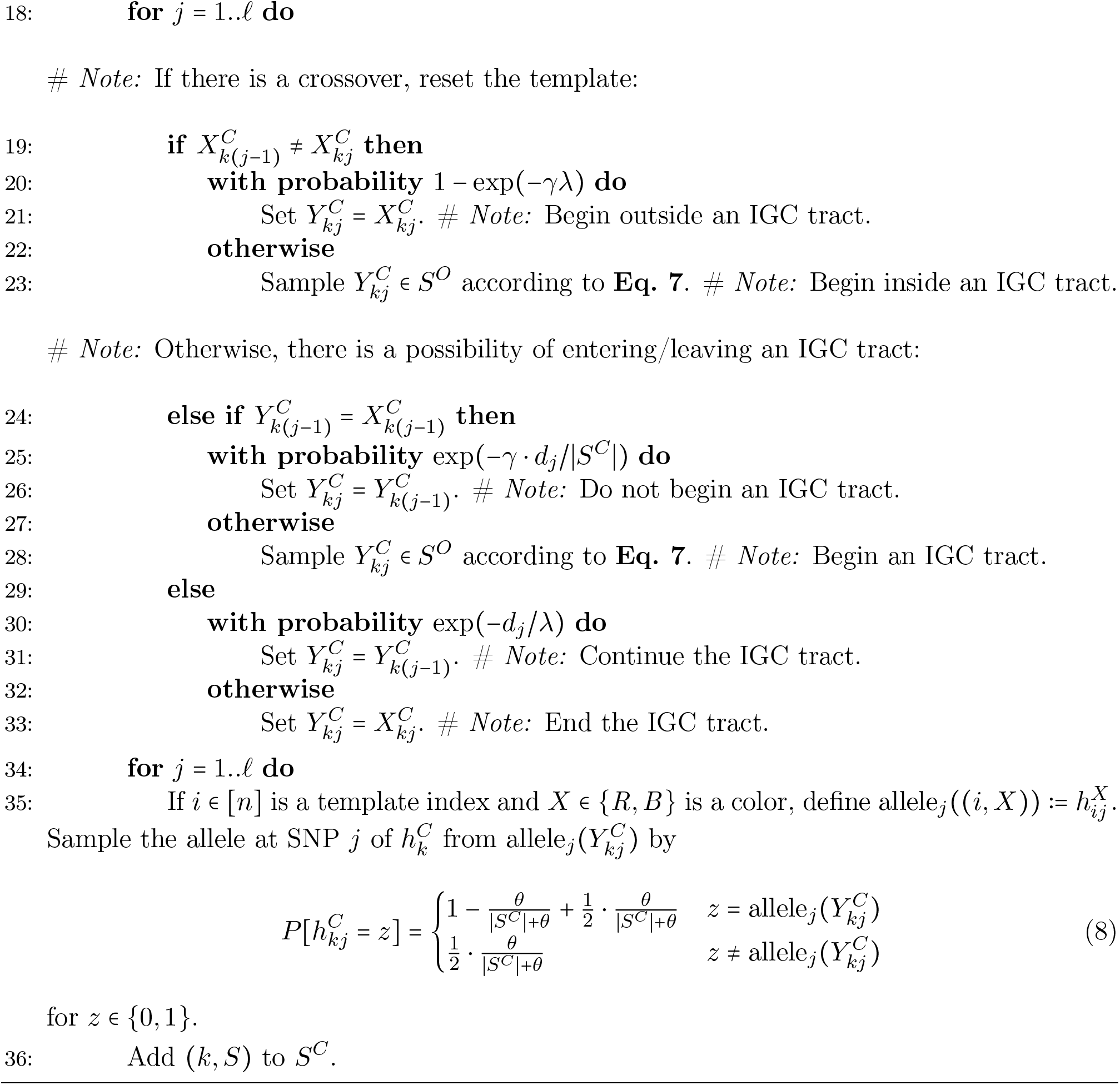

## 3 Validation

In this section, we discuss the results of a simulation study evaluating the utility of LS4Dups in estimating IGC parameters and for detecting IGC havens. Before presenting the simulations and their results, we outline qualitative and quantitative goalposts we set for LS4Dups. In the standard LS model [33], a parameter *ρ* governs the frequency of switches between copying templates and serves as an proxy for the molecular crossover rate, *ρ*^∗^. *ρ*^∗^ and copying model parameter *ρ* are not exactly equivalent, because *ρ* is based on a heuristic model of the TMRCA between templates and foci. However, a useful estimator for the molecular rate, 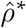 has been proposed by [33] based on assuming the relationship

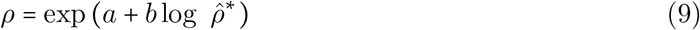

with *a* and *b* fitted in a standard generalized linear regression model.^3^

We likewise do not expect that the IGC initiation and mean tract length parameters in LS4Dups will be equal to the molecular parameters we wish to estimate. For LS4Dups to be useful in the study of the molecular parameters of IGC and their genomic variation, it would suffice for the relationship between inferred LS4Dups parameters and molecular pararameters to prove to be monotonic—or ideally near-linear on a log scale—as in [33].

We tested LS4Dups’s utility in measuring IGC parameters, the initiation rate and the mean tract length, on simulated data. We also tested its ability to infer the population-scaled per-site IGC rate, meaning the probability of a particular site being in an IGC tract over 4*N* generations. We estimated the parameters of LS4Dups representing initiation rate and mean tract length, respectively *γ* and *λ*, by maximum likelihood inference on the simulated data. Our simulations are intended as a proof of concept for LS4Dups and its ability to measure IGC rate under ideal conditions, and our simulation parameters are not chosen as biologically realistic values for humans or any particular organism, nor do we attempt to model the effects of demography or population size change.

For simplicity, we assumed the LS4Dups crossover parameter *ρ* to be known and equal to the generative crossover rate. The reason we did not infer crossover rate directly is that in experiments we observed that including crossover rate as a variable reduces the precision of IGC parameter estimates. In future application to real data, one possible approach may be to infer LS4Dups *ρ* by an LS-based *ρ* estimate from a flanking, non-duplicated sequence.

Our simulations were run in SLiM [48]. We describe the simulation methods and parameters in more detail in **Supplementary Note Section S1**. Each simulation (**Fig. 5a**) began with a population of 2*N* haplotypes that included two duplicate loci of length 1 MB, with a 1 MB region between them. The simulation was divided into two periods. The first was a “burn-in” period during which there was crossover and mutation, but no IGC, increasing the baseline polymorphism levels within the two duplicates to approximate the process of the duplication reaching fixation and some polymorphism being created between and among the two regions.

**Figure 5.**
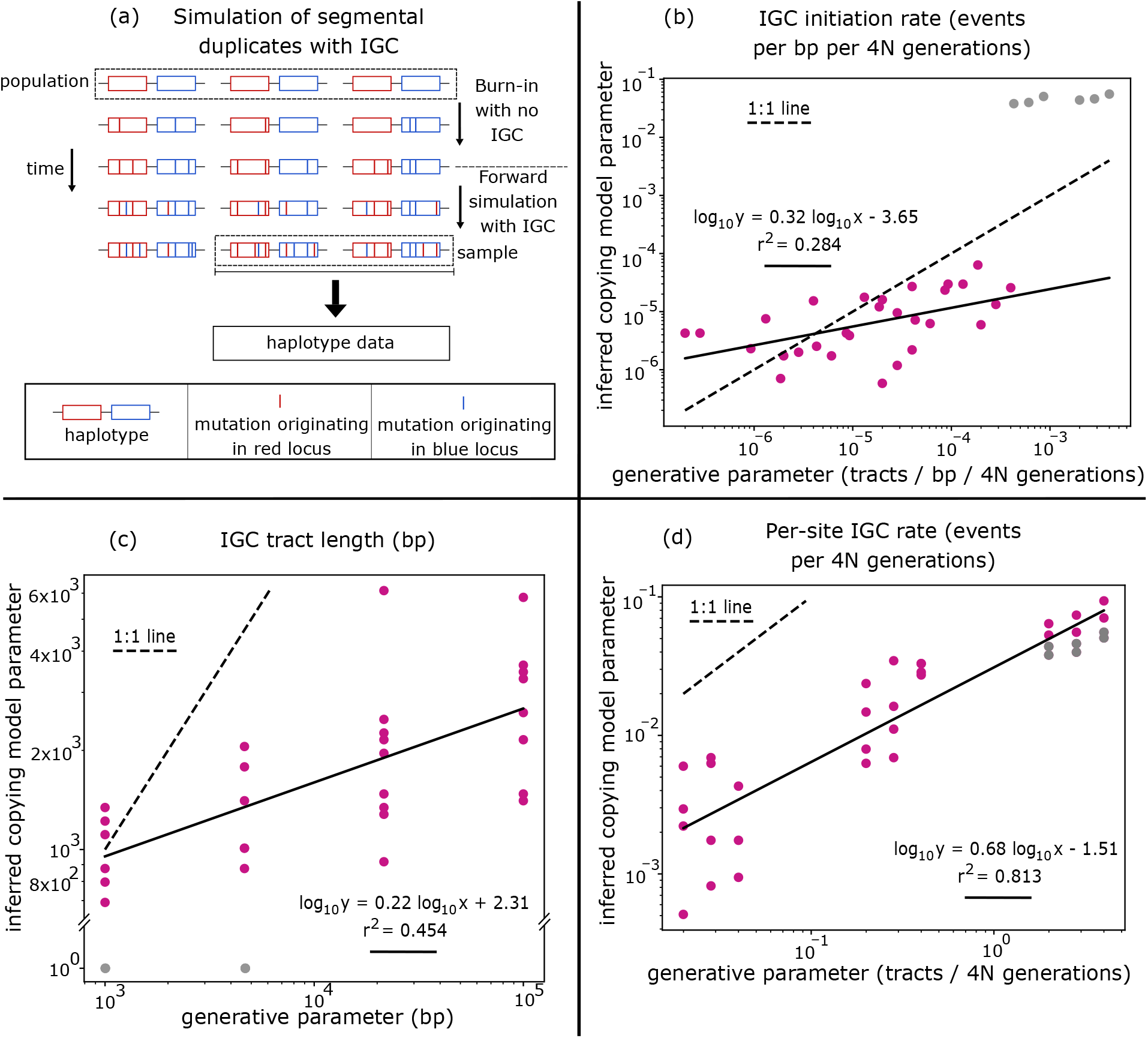
Parameter inference on simulation. **(a)** Schematic of our forward-in-time Wright-Fisher simulation of the evolution of a population of segmental duplicates. The simulations start with a period with mutation and crossover but no IGC, meant to generate initial polymorphism patterns, and followed by a period with IGC between the duplicates. **(b-d)** Shown are maximum likelihood estimates from the application of LS4Dups to 34 simulations. LS4Dups copying model parameters (y-axes in **(b**,**c)**) do not correspond exactly to the generative parameters (x-axes in **(b**,**c)**), but they track them monotonically. **(d)** shows that for the per-site IGC rate, the product of initiation rate and expected tract length, the relationship between copying model parameter and generative parameter is tighter than for these parameters individually. Each point corresponds to an approximate maximum likelihood estimates based on a grid search in one iteration of the simulation. Simulations in which the maximum likelihood estimate was at at its minimum value 1) are shown in grey. In **(b)** and **(c)**, grey points are qualitatively far outside the variation pattern of the other points, and are omitted from the OLS regression in **(b)** and **(c).** In **(d)**, grey points are qualitatively within the varation pattern of the other points, and are included in the OLS regression. Shown with black solid lines are OLS fits on a log-log scale. Black dashed lines indicate the one-to-one line.

The second period introduced IGC into the simulation, and ran for 2*T* generations^4^ where *T* is the expected TMRCA of the full population.

We applied maximum likelihood estimation using a grid search to the simulated data (**Fig. 5b-d**). The LS4Dups IGC initiation rate 4*Nγ* (**Fig. 5b**) and mean tract length parameter *λ* (**Fig. 5c**) correlated weakly with the corresponding molecular values (*r*^2^ = 0.284 and *r*^2^ = 0.454 respectively). The relationship of both these inferred model parameters to the generative parameters was close to linear on a log-log scale but noisy, and with several points at high IGC initiation rate and low mean tract length appearing to be qualitative “outliers” (shown in grey). However, the estimated overall per-site IGC rate across the full duplicated region, which is the product of these two, was more strongly correlated with the generative parameter (*r*^2^ = 0.813), with a close-to-linear relationship between the two on a log-log scale, and the same qualitative outlier points falling close to the trendline (**Fig. 5d**).

### 3.1 Detecting IGC havens as signatures of selection favoring differentiation of duplicates

Paralogous sites may be selected for different alleles. For example, the X-linked cone opsin genes in humans have particular sites which differentiate the wavelength of light detected by the opsins [9]. In such a case, the site selected for divergence, as well as strongly linked sites, are shielded from IGC’s homogenization of alleles at the paralogous sites. Even though LS4Dups (like other ARG inference methods) inherently assumes neutral evolution, we would expect lower in-sample evidence of IGC and, consequentially, a decrease in inferred IGC rates close to the selected site. We call this hypothesized signature an *IGC haven* after the expected “shielding” reducing inferred IGC rate. We further expect the width of the IGC haven to scale with the mean tract length parameter, because for a site at fixed distance from the selected site, a longer mean tract length implies that the site occurs in IGC tracts along with the selected site more frequently, and therefore fewer IGC tracts at this site survive selective purging. For the signature at linked sites to be detectable, the haven must reduce the inferred IGC rate to below the inferred levels of IGC in flanking sequence, and this dip must be larger than the background variation in inferred IGC rate. We expect the depth of the reduction to increase with stronger selection, and the background variation in inferred IGC rate to decrease with stronger IGC (**Fig. 5d**). Therefore, only only under both strong IGC and strong selection for divergence do we expect to see IGC havens (**Fig. 6a**).

**Figure 6.**
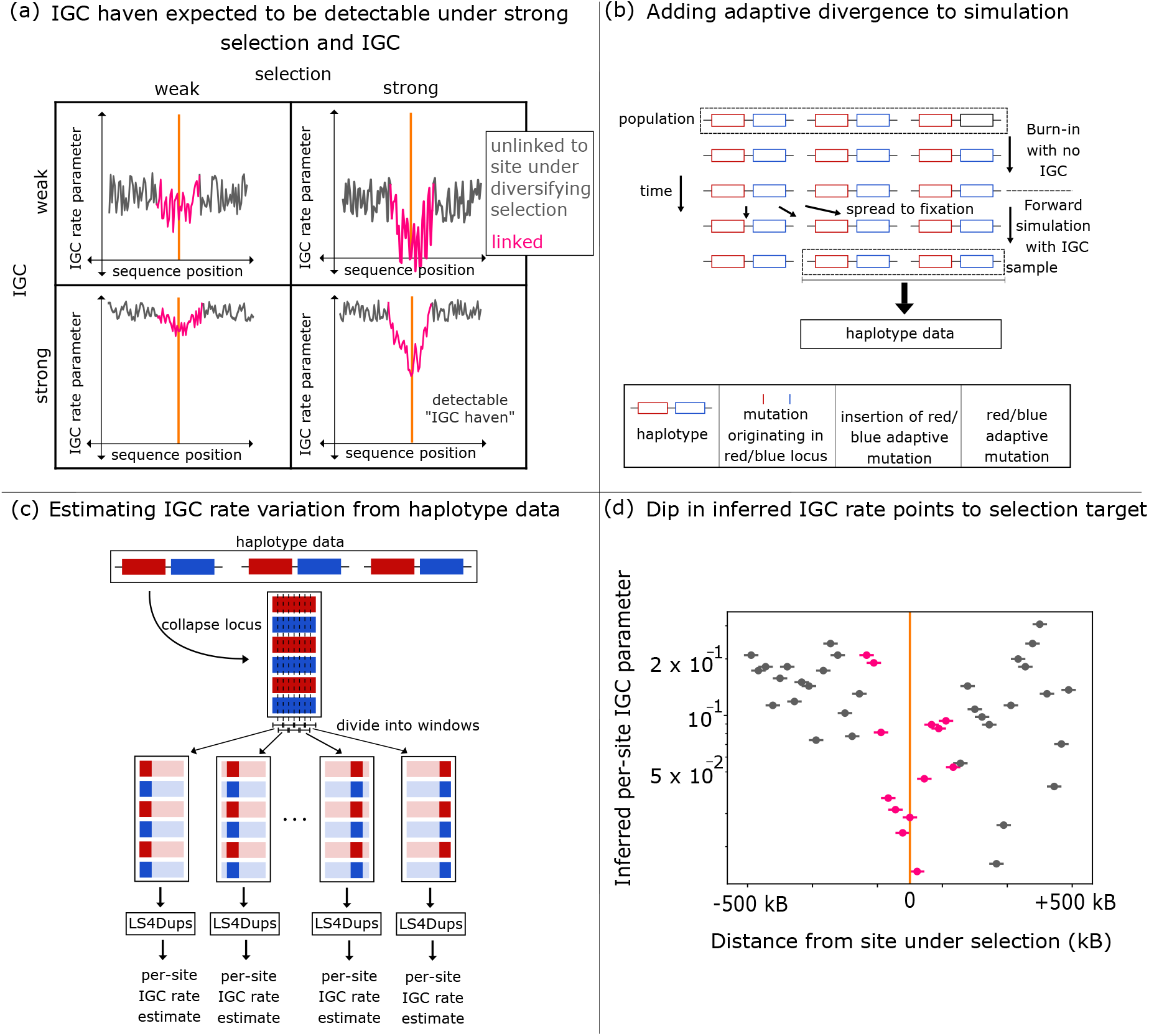
Detection of IGC haven. **(a)** Cartoon example of the conditions under which we expect the IGC haven (reduction in IGC due to selection for sequence divergence) to be large enough relative to background variation to be detectable: strong selection and strong IGC. The orange vertical line shows the site under selection for divergence between the duplicates. **(b)** Modification of the simulation in **Fig. 5a.** Following the burn-in period, an adaptive divergence was introduced in a randomly selected chromosome. The two introduced alleles were positively selected when in their matching-color locus, and selected against when in the other-color locus. **(c)** We applied LS4Dups in a sliding window of fixed width. **(d)** Per-site IGC rate prediction in one simulation (out of 10) with an inserted divergent site under strong selection for divergence (orange line). Points represent the centers of windows, and horizontal bars indicate the extent of the window over which inference was performed. Pink points are those windows whose centers are within 100 kB (twice the mean IGC tract length) of the site under selection for divergence. The IGC haven signature, a sharp spike downward in IGC rate, was visible near the selected site in some of the simulations including the one shown, but not all. The pink points in the trial shown diverge significantly from the grey points under a two-sample T-test (alternate windows overall divergence of sites within 100 kB of the selected site was also significant (*T* = 5.589, *p* < 10^−7^).

To illustrate the ability of LS4Dups to detect IGC havens in such a parametric regime, we added an adaptive divergence at a single site to our simulation (**Fig. 6b**). Namely, at the end of the burn-in period, a new “red allele” was introduced at a site in the center of the red locus, and a “blue allele” in the center of the blue locus. The fitness function was defined based on a selection strength parameter *s*, so that the red allele had a selective coefficient of *s* in the red locus and −*s* in the blue locus, and the blue allele had a selective coefficient of *s* in the blue locus and −*s* in the red locus.

In our inference, we conditioned the simulation on fixation of the adaptative divergence by restarting the simulation after the burn-in with a newly sampled random seed when the new adaptations were lost. We intentionally chose a set of parameters with high IGC rate, large IGC tracts, high mutation rate, and strong selection to demonstrate the potential of the method.

For each simulation, we broke the collapsed locus into overlapping windows of fixed width 50 kB (**Fig. 6c**). We ran LS4Dups maximum likelihood inference on each window independently. In our simulations, the region near the selected site differed significantly from the rest of the collapsed locus in inferred IGC rate (*p* < 10^−7^), although results were noisy (**Fig. 6d** and **Supplemental Fig. S1**).

## 4 Discussion

Following coalescent models incorporating IGC developed by Innan [31] and Thornton [32], we described the Collapsed Site Coalescent, a representation of the genealogical process at a pair of segmental duplicates that is suited to ARG inference. We used the CSC to build LS4Dups, an adaptation of the LS model that estimates IGC parameters from a population sample of haplotypes. In simulations, LS4Dups estimates track the per-site IGC rate, and reveal IGC havens around sites under selection for divergence between duplicates when both IGC and selection are strong.

The promise of this approach lies in the information contained in polymorphism data. For point mutation and recombination, population genetic analyses of polymorphism have transformed what can be learned, compared with the two other main sources of data. Divergence between species averages over millions of years and cannot resolve fine-scale or species-specific variation; recombination hotspots, for example, are largely not shared between humans and chimpanzees [49, 50]. Experiments and family studies observe individual events directly [13, 51–53], but over only a few generations, and so see few events at any given position. Polymorphism data instead record the events along the genealogy of a sample back to its most recent common ancestor—many more generations—while still reflecting recent, species-specific features of the molecular processes involved. Fine-scale recombination maps inferred from linkage disequilibrium with the LS model and related methods showed that recombination is concentrated in narrow hotspots and pointed to their determinants [33, 54–56]: PRDM9, which positions meiotic double-strand breaks in humans and mice [57, 58], and, in the several vertebrate lineages in which PRDM9 has been lost wholly or in part, hotspots at promoters and CpG islands, which can persist over long evolutionary timescales [59–61]. Likewise, polymorphism in large population samples revealed the extent of mutation rate variation along the genome and its sequence-context drivers [62–64], underpinned measures of mutational constraint on genes [65, 66], and showed that the mutation spectrum evolves rapidly among human populations and across species [67– 69], far beyond what family and divergence data had resolved.

IGC has so far been studied mostly through divergence between species, family studies, or a small number of loci [14, 70, 71], and exploiting polymorphism data could similarly transform the study of IGC. Full exploitation of polymorphism data requires sequencing that can resolve the alleles carried by each paralog, which short reads cannot provide in highly similar duplicates. omitted to remove overlap, *T* = 3.415, *p* < 0.0015). Out of 10 trials (**Supplementary Fig. S1**), the Population samples of haplotype-resolved, long-read assemblies are now available in humans [72, 73] and have already been used to characterize single-nucleotide variation, IGC, and structural variation in segmental duplications [3, 15]; similar resources are accumulating in other species [74]. LS4Dups, and its extension to scalable ARG inference, are designed for such data.

Whether polymorphism data contain enough signal to estimate IGC parameters and their variation accurately, and to detect IGC havens, in real segmental duplicates is an empirical question. We expect the signal to be strongest where background polymorphism is high, IGC is frequent, and selection for divergence is strong.

Our simulations used a simple scenario, a single site under symmetric selection for divergence with frequent IGC and long conversion tracts, whereas adaptive divergence between duplicates can involve multiple interacting sites, as in the one case we have studied empirically [8]. A more comprehensive investigation of possible modes of selection, as well as demographic history, the age of the duplication, and variation in IGC and crossover rates will all shape the signal and its detectibility. These effects remain to be characterized.

Because the LS model is the core of several scalable ARG inference methods [23, 24], a natural next step to our work is to extend LS4Dups to infer full ARGs for segmental duplicates. We outline how this could be done within the framework of Relate [23] in **Supplementary Note Section S2**. The main new challenges are that whether two paralogous lineages are carried on the same chromosome must be inferred indirectly, and that short IGC tracts make genealogies change more frequently along the sequence, which complicates matching branches between neighboring genealogies.

Finally, we considered pairs of duplicates with a fixed copy number. The CSC extends in principle to larger gene families of fixed size, at the cost of a larger set of lineage types, but assumes that copy number does not vary within the genealogical history of the sample. This is also true of Innan’s coalescent model [31] but not Thornton’s [32], which allows for gene duplication to occur within the genealogy of the sample. Variation in copy number among haplotypes is common in human segmental duplications [3], and incorporating it into genealogical inference would require a different representation and remains an important challenge for future work.

Segmental duplicates make up a substantial fraction of genomes and are a major source of evolutionary novelty; yet, they have remained largely outside the reach of the genealogical methods that have transformed population genetics elsewhere in the genome. By making genealogical inference possible in the presence of IGC, the CSC and LS4Dups open the way to mapping IGC rates across segmental duplications, relating their variation to molecular features such as sequence identity, distance between paralogs, and G/C content and quantifying how IGC shapes the evolution of duplicates. Our models also lay the groundwork for scanning population samples for IGC havens, which, when IGC and selection are strong, can pinpoint the specific sites at which selection has maintained functional differences between duplicates. As haplotype-resolved population samples accumulate across species, these tools could help reveal how duplicate sequences evolve and the conditions that allow them to functionally differentiate.

## Footnotes

1 Throughout we assume a segmental duplication that has fixed, so that the copy number is constant within the population

2 Although in order for the expected number of mutations per SNP to be 1 as in the LS model, we should instead choose the reciprocal of the TSBL described in Section 2.1, we found that a value of *θ* which depended on the crossover and IGC rates interfered with inference accuracy. In the classic LS model, *θ* is a parameter independent of crossover rate, and we preserve this independence from recombination rates in our implementation.

3 In the original work this is presented as a multiplicative correction term *ξ* = exp(*a* + *b* log *ρ*^∗^) such that the model crossover rate is *ρ* = *ρ*^∗^*ξ*, but our formulation is mathematically equivalent.

4 We kept this period short due to computational constraints—our implementation of IGC in SLiM slowed it down substantially. To ensure that mutations are realistically distributed, there should be a single most recent common ancestor of the full sample (the tree+ root for the sample) within the simulation at every collapsed site, and a collapsed site is more likely to have a single most recent common ancestor the more generations the simulation is run for.

## Notes

### Competing Interest Statement

The authors have declared no competing interest.

