## Supplemental Note for "Collapsed Site Coalescent Facilitates Detection of Adaptive Divergences Between Segmental Duplicates"

### Methods Supplement

October 23, 2026

#### Contents

**S1 Simulation Details** **1**

**S2 Considerations for Full ARG Inference** **2**

**S3 TMRCA and TSBL Computation** **8**

S3.1 Approximate computation . . . . . . . . . . . . . . . . . . . . . . . . . . . . . . . . 8

S3.2 Exact computation . . . . . . . . . . . . . . . . . . . . . . . . . . . . . . . . . . . . 8

### S1 Simulation Details

Simulations are run in SLiM [1], with a 3 MB region divided into three parts: two 1 MB segmental duplicates flanking a 1 MB sequence between them. We introduce IGC into SLiM by sampling breakpoints after every generation, sampling tract endpoints by a geometric distribution of tract length (truncating tracts to stay within the 1 MB region), and manually creating new mutations at the equivalent site in the paralogous locus to unify the mutation pattern between the two duplicates within the tract for the affected chromosome. At high per-site IGC rates the creation of so many new mutation objects significantly slows down SLiM, but it was necessary to implement this way rather than treating duplicates as separate samples in a population in order to capture the coupling property of the duplicates. The simulation begins with a period of burn-in during which mutation and crossover occur but no IGC. The mutation rate is set at a higher value before the burn-in, to generate additional background polymorphism. Immediately after the burn-in period, IGC begins to occur, and the mutation rate is reduced.

In the simulations involving selection, immediately after the burn-in is also when we add the selectively-maintained divergent site. It is introduced as a special type of mutation, one in each duplicate, with a positive selective coefficient $s$ occurring in the exact middle of each duplicate, such that if the mutation originating in the "red" duplicate is copied into the "blue" duplicate, its selective coefficient becomes $-s$. A single haplotype is chosen at random from the population and the corresponding mutation is added to each of its duplicates. The program saves its state before continuing with the post-burn-in generations, and if after any generation one or other of the selected mutations is not present in the population, the simulation is reset immediately at the end of the burn-in with a newly sampled random seed. This ensures that the divergence fixes in the population.

In our experiments estimating the rates of IGC over varying generative rates (main text **Fig. 5**), we set the burn-in mutation rate to $10^{-6}$ mutations per base pair per generation, the mutation rate after burn-in to $10^{-7}$ mutations per base pair per generation, and the crossover rate to $10^{-8}$ crossovers per base pair per generation. We varied other parameters in even increments on a log scale: population size in 5 values between 500 and 2000 diploid individuals, per-site IGC rate in 4 values between $10^{-3}$ and $10^{-6}$ gene conversions per site per generation, and mean IGC tract length in 4 values between 1 kB and 100 kB. Of the 80 combinations of these parameters, many with high population size and high per-site IGC rate were omitted due to time constraints; we ran 34 simulations total. The burn-in lasted for $20N$ generations with $N$ the diploid population size, and the number of generations after burn-in was twice the expected TMRCA, approximated as described in **Section S3**.

In our experiment detecting the IGC haven selective signature (main text **Fig. 6**), the mutation rate was $10^{-6}$ mutations per base pair per generation before burn-in and $10^{-7}$ after, and the crossover rate to $10^{-8}$ crossovers per base pair per generation. The population size was 200 diploid individuals, the per-site IGC rate was $2 \times 10^{-2}$ gene conversions per site per generation, the mean tract length was 50 kB, and the selective coefficient $s$ for the divergent adaptation was 0.9. The burn-in lasted $20N = 4,000$ generations, and the number of generations after burn-in was $2N = 400$.

For inference in both experiments, we sampled 12 haplotypes and used LS4Dups to infer the likelihood of generating the haplotypes given IGC initiation and mean tract length model parameters. We used a grid search to find the maximum-likelihood parameter combination, with a $9 \times 9$ grid, iterated 5 times, with each iteration defining a new grid with $2/9\times$ the lattice width, centered on the highest likelihood point in the previous grid for the next iteration. The initial grid varied from $10^{-10}$ to 1 for IGC initiation, and from 1 to $10^{10}$ for mean tract length. The full collapsed locus was used for the rate estimation experiment. For the IGC haven experiment, we divided the 1 MB collapsed duplicated region into $W = 45$ windows of length $2/(W+2)$ MB, with the $i$th window beginning at $i/(W+2)$ MB ($i = 0, \ldots, (W-1)$), and each window was given to LS4Dups separately.

#### S2 Considerations for Full ARG Inference

The classic Li-Stephens (LS) model is the basis of several methods for inferring a full Ancestral Recombination Graph (ARG), including Relate [2] and tsinfer [3]. In the case of Relate, the LS model is repurposed to estimate the dissimilarity (the amount of divergence) between each pair of samples at each site, forming an asymmetric dissimilarity matrix $D$ between the samples. Here $D(a, b)$ is an estimate of the number of alleles derived in $a$ and ancestral in $b$, or equivalently (in an infinite site model) the number of mutations occuring on the branch from $a$ to its most recent common ancestor with $b$. $D(a, b)$ is computed by a linear translation from the log probability that $b$ is the template $a$ copies from in an LS Model with every pseudo-sample but $a$ available as a template. This value is chosen so that for $b, b' \neq a$, in the limit as recombination rate goes to zero, $D(a, b) < D(a, b')$ exactly when $b$ coalesces with $a$ sooner than $b'$. This approximates

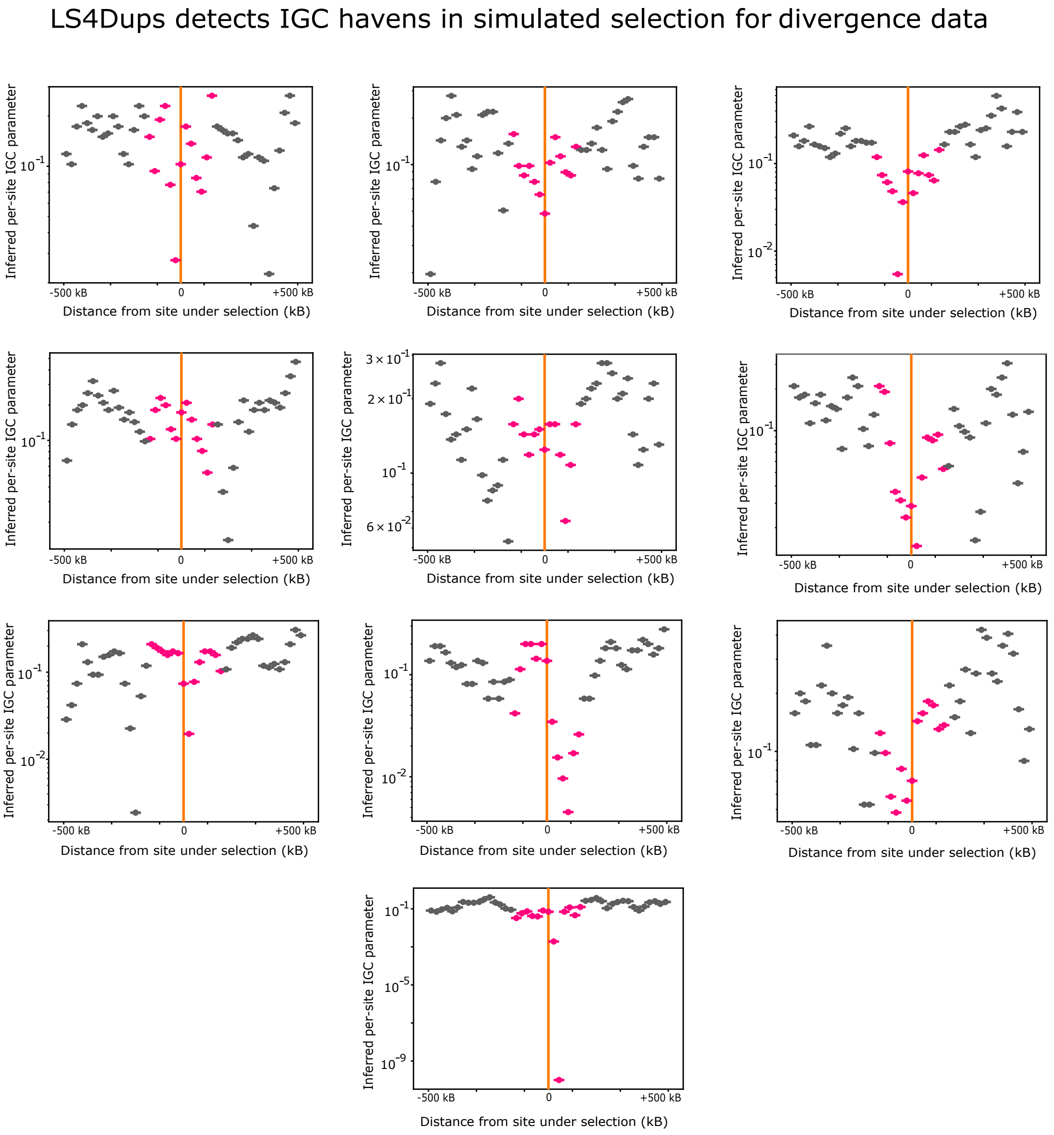


**Supplemental Figure S1. Simulations of IGC Havens**. Results for 10 simulations of selection for divergence. Pink points represent points within twice the mean tract length (100 kB) of the divergence site. The simulation procedure is illustrated in main text **Fig. 6**. The inferred values of IGC rate for sites within 100 kB of the selected site were signficantly different from the other values (omitting alternate windows so that windows did not overlap, $T = 5.589$, $p < 10^{-7}$.)

the true ordering of the neighbors of each sample. This matrix is used to reconstruct a tree topology (not including the branch lengths) at each site, through a form of clustering closely based on UPGMA [4], with optimizations to allow for reuse of estimated topologies when possible. UPGMA (unweighted pair group method with arithmetic mean) begins with all elements in their own singleton sets. The distance between sets $\mathcal{A}$ and $\mathcal{B}$ is computed by

$$D(\mathcal{A}, \mathcal{B}) = \frac{1}{|\mathcal{A}| \cdot |\mathcal{B}|} \sum_{a \in \mathcal{A}} \sum_{b \in \mathcal{B}} D(a, b). \quad (1)$$

The algorithm will join pairs of sets $(\mathcal{A}, \mathcal{B})$ such that each is the most similar to each other, that is, $D(\mathcal{A}, \mathcal{B}) \leq D(\mathcal{A}, \mathcal{B}')$ for all $\mathcal{B}' \neq \mathcal{A}$ and $D(\mathcal{B}, \mathcal{A}) \leq D(\mathcal{B}, \mathcal{A}')$ for all $\mathcal{A}' \neq \mathcal{B}$ (ties broken by symmetrized distance $D(\mathcal{A}, \mathcal{B}) + D(\mathcal{B}, \mathcal{A})$). The selected pair is then joined into a new set $\mathcal{A}' = \mathcal{A} \cup \mathcal{B}$, with the join recorded as a coalescence in the tree, so this method builds the tree topology. To estimate branch lengths, because many ARG branches occur in multiple adjacent trees, it is necessary to identify which tree branches in adjacent trees correspond to the same ARG branch. Pairs of tree branches to be matched are identified by the descent patterns of their parent and child nodes. An overall mutation rate for each ARG branch can then be estimated, which is used as part of an MCMC model to estimate the branch lengths of the trees.

A natural extension of our adaptation of the LS model would be to inference of full ARGs for duplicated genes, using the Relate methodology. We note here several considerations that would be involved in making this adaptation. First considering the construction of the pairwise dissimilarity matrix, our adapted HMM may be used to generate a dissimilarity matrix in the same way as in Relate (via the same linear transformation), but the inclusion of IGC complicates the interpretation of the matrix. A naïve use of UPGMA clustering on collapsed locus data is susceptible to errors such as the one in **Supplemental Fig. S2**. In this example, the pseudo-samples red 1, blue 1, and red 2 all form a clade sharing a derived mutation at SNP 2. In the true tree, this is due to an IGC event occurring on the branch from leaf 1 to the coalescence of leaves 1 and 2, which spread the mutation from red to blue. Although there is no mutation to indicate that red 2 is the outgroup, placing red 2 as the outgroup requires only an IGC event to have occurred, while any other arrangement requires at least one decoupling in addition, and so is less parsimonious. However, because red 1's sequence can be generated from red 2 as a template under our adapted LS model without an IGC event being needed, $D(\text{red } 1, \text{red } 2) < D(\text{red } 1, \text{blue } 1)$, so UPGMA would join red 1 to red 2 before blue 1. This type of error could be corrected by changing the clustering algorithm to include an additional parameter $\xi$ punishing joins that require a decoupling. In the example, we would instead compare $D(\text{red } 1, \text{red } 2) + \xi$ and $D(\text{red } 1, \text{blue } 1)$. It may also be possible to address this problem by a adjusting the translation from the HMM probabilities to the dissimilarity matrix.

To adapt Relate's joining of branches in adjacent trees (for mutation counting and estimation of branch lengths), we must join branches in adjacent tree+s. Adjacent tree+ topologies may differ from each other in more extreme and less localized ways than adjacent tree topologies. **Supplemental Fig. S3** shows one example in which a decoupling and a branch associated with it disappear due to the start of an IGC tract several branches down on the tree, and then reappear due to the end of a different IGC tract. In terms of the true full ARG, this is a single branch, and fully capturing all available information on its mutation rate requires modeling that it is a single branch that is present in two tree+s but not present in a tree+ between them, a

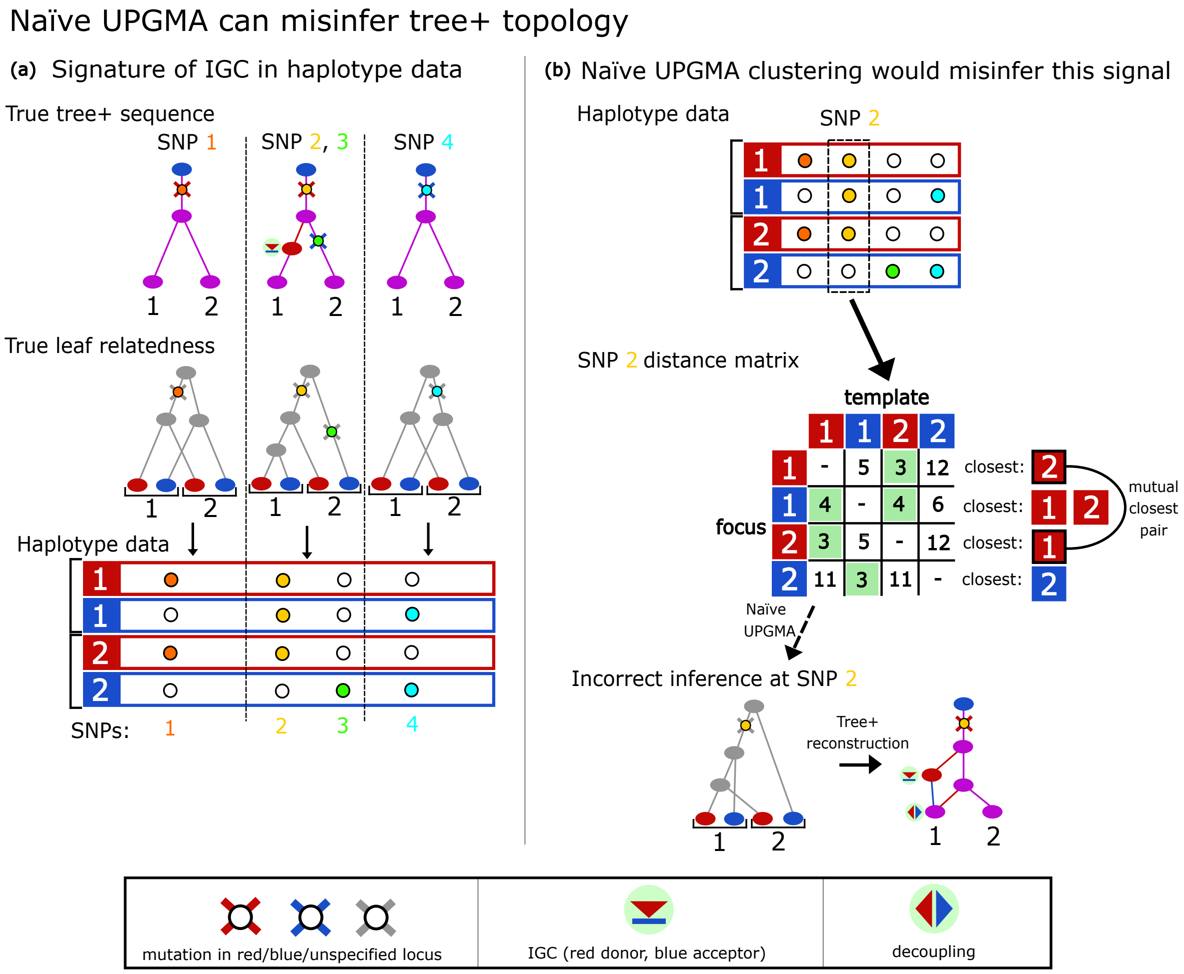


**Supplemental Figure S2. Complications in tree+ topology inference.** **(a)** An example of a short 4-SNP sequence generated by 3 adjacent tree+, including an IGC event covering SNPs 2 and 3 and affecting sample 1. This IGC introduces the derived mutation at SNP 1, which originated in the red locus, into the blue locus. **(b)** A hypothetical inference of the tree+ topology at SNP 2. The numbers used in the dissimilarity matrix are made up for the point of illustration, but their relative magnitudes are consistent with the data shown. Notably, the dissimilarity from red 1 to red 2 is smaller than the dissimilarity from red 1 to blue 1, since copying from red 2 does not require an IGC event to have occurred. Naïve UPGMA clustering on this matrix would join red 1 and red 2 first as they are each other's nearest neighbor, which requires a decoupling not present in the true tree+. One possible tree+ reconstruction is shown; others are possible, but also require more events to explain.

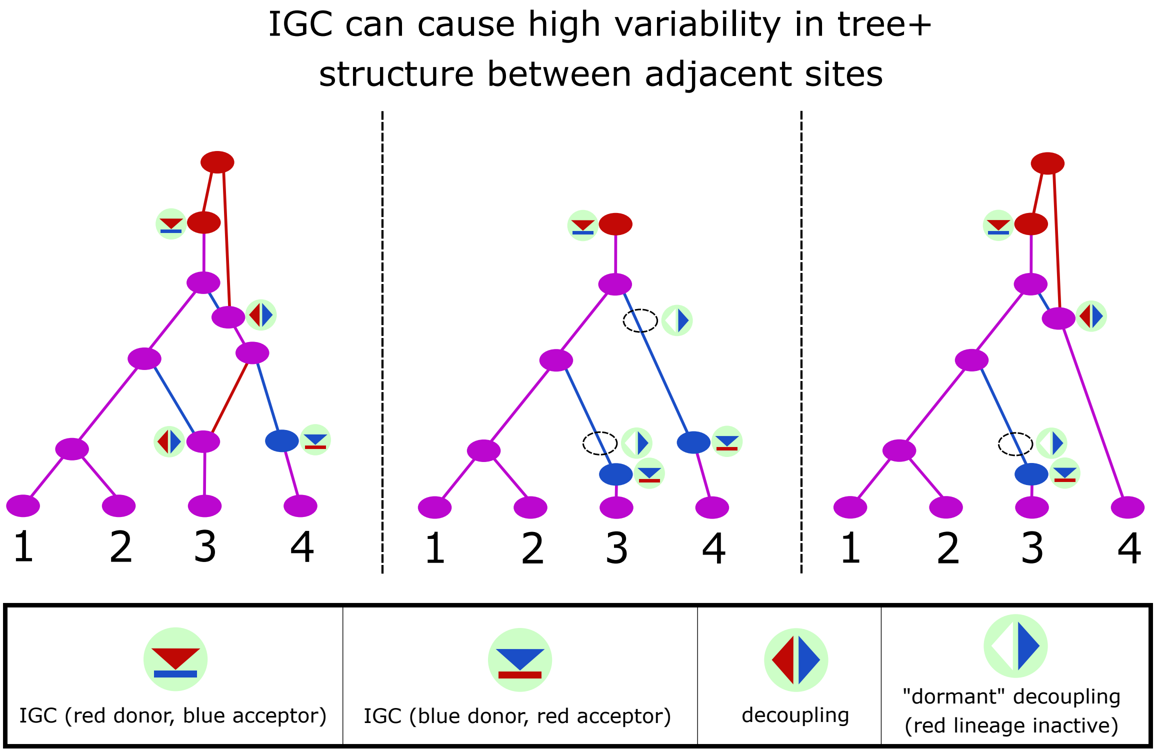


**Supplemental Figure S3. Complications in joining branches of adjacent tree+s.** A series of three adjacent tree+s. The first tree+ has two red branches at its top which coalesce at the root. In the second tree+, the IGC event immediately above leaf 3 causes two decoupling events to become inactive, and removes these red branches from the tree+. In the third tree+, another IGC above leaf 4 ends, which reactivates the higher decoupling and reintroduces the same red branches back into the tree+.

"non-Markovian gap." With high IGC rate, such non-Markovian gaps may be very common, and if branches are only joined across adjacent trees, this may lead to fragmentation that reduces the accuracy of branch length estimation. A branch joining method for tree+s may therefore need to involve comparing branches in non-adjacent trees, possibly trees within some fixed number of base pairs of the current tree. Since branches are identified as equivalent based on descent pattern, it will also be necessary to account for the fact that descent pattern can shift rapidly due to IGC — a node may gain or lose many descendants of a new color if a new IGC occurs below it.

### S3 TMRCA and TSBL Computation

For various purposes, including setting the interal parameters of the LS-style HMM, it is useful to compute the expected Time to Most Recent Common Ancestor (TMRCA) and expected Total Sub-Branch Length (TSBL) for the tree+. In the standard coalescent, with no decoupling or conversion, these questions admit considerable mathematical simplification — the expected TMRCA of the tree (in coalescent units) reduces to $\sum_{k=2}^{n} 1/\binom{k}{2}$ which simplifies to $2(1-\frac{1}{n})$. The expected TSBL of the tree reduces to $\sum_{k=2}^{n} k/\binom{k}{2}$ which is a harmonic sum well approximated by $2\log n$. Underlying these simplifications is a desirable property of the standard coalescent: that the coalescent rate parameter depends only on the number of existing lineages, which follows a deterministic trajectory from $n$ to 1. Whenever there are $k$ lineages, the rate parameter to the next event is always $\binom{k}{2}$, and the next state is always $k-1$ lineages.

#### S3.1 Approximate computation

In the CSC these conditions do not hold. Defining the *generation sub-branch count (GSBL* as the number of active sub-branches at a particular time, the GSBL does follow a deterministic trajectory from $2n$ to 1. However, there are many possible configurations of GSBL $k$, which give rise to differing event rates. The expected time to move from GSBL $k$ to $k-1$ is dependent on the initial lineage configuration. It is possible to exactly compute the expected TMRCA and TSBL using $O(n)$ matrix multiplications of size $O(n^2) \times O(n^2)$ using a dynamic programming method described in section S3.2. Depending on the form matrix multiplication used, this could take approximately $O(n^6)$ time.

To avoid the high cost of this exact inference, we introduce an approximation, based on the following observation: In the absence of IGC, each of the two sites will behave like a standard coalescent tree, and the expected time until the final red coalescence is $2(1-\frac{1}{n})$ as in the standard model, and the same holds for the blue coalescence. The introduction of IGC will introduce noise into the time until final coalescence of each tree, since the number of extant lineages may now be increased or decreased by an IGC, but this affect is balanced such that the expected net flow of lineages between the A and B coalescent trees is zero. Therefore we expect that the effect of IGC on the coalescence time will only be appreciable when two lineages remain. We may use the exact model below to compute the expected time $T_2(R,C)$ to final coalescence of two lineages, assuming they are coupled with probability $\frac{1}{1+R}$ and decoupled with probability $\frac{R}{1+R}$ ("coupling-decoupling balance"). We therefore approximate the TMRCA by $2(1-1/n)+T_2(R,C)$. We may likewise approximate the TSBL by $4\log n + 2\cdot T_2(R,C)$.

#### S3.2 Exact computation

The set of configurations of GSBL at most $k$ is given by all triples of whole numbers representing the number of lineages in each state (purple, red, or blue):

$$S_k := \left\{(n_P, n_R, n_B) : {}^{n_P \geq 0,\, n_R \geq 0,\, n_B \geq 0,}_{2n_P + n_R + n_B = k}\right\} \subset \mathbb{Z}^3. \tag{2}$$

For $k > 1$, represent the expected TMRCA and TSBL when the starting state is some state $s \in S_k$ by $d(s)$ and $w(s)$ respectively. We will end up computing vectors $\mathbf{d}_k$ and $\mathbf{w}_k$ representing the

| event type | new arrangement | rate |
|---|---|---|
| purple-purple coalescence | $(n_P - 1, n_R, n_B)$ | $\binom{n_P}{2}$ |
| purple-red coalescence | $(n_P, n_R - 1, n_B)$ | $n_P n_R$ |
| purple-blue coalescence | $(n_P, n_R, n_B - 1)$ | $n_P n_B$ |
| red-red coalescence | $(n_P, n_R - 1, n_B)$ | $\binom{n_R}{2}$ |
| blue-blue coalescence | $(n_P, n_R, n_B - 1)$ | $\binom{n_B}{2}$ |
| coupling | $(n_P + 1, n_R - 1, n_B - 1)$ | $n_R n_B$ |
| decoupling | $(n_P - 1, n_R + 1, n_B + 1)$ | $R n_P$ |
| purple-to-red color change | $(n_P - 1, n_R + 1, n_B)$ | $C n_P$ |
| purple-to-blue color change | $(n_P - 1, n_R, n_B + 1)$ | $C n_P$ |
| red-to-blue color change | $(n_P, n_R - 1, n_B + 1)$ | $C n_R$ |
| blue-to-red color change | $(n_P, n_R + 1, n_B - 1)$ | $C n_B$ |

**Supplemental Table S1.** Table of possible transitions from a lineage arrangement $(n_P, n_R, n_B)$ representing the number of purple, red, and blue lineages respectively. The waiting time to the next event may be modeled as an exponential distribution with rate parameter given by the sum of the rates of the possible next events, where the probability of a particular event being the next is proportional to its individual rate contribution. The rates are scaled in standard coalescent units such that the expected time for a pair of lineages to coalesce is 1. $R$ and $C$ are respectively the decoupling and conversion parameters of the tree+.

expected values of TMRCA and TSBL for every $s \in S_k$. After some period of time (possibly involving transitions between different GSBL $k$ states) the GSBL will transition to a value below $k$[1]. Let $T_k \subseteq S_k \cup S_{k-1} \cup S_{k-2}$ be the set of states which can be transitioned to directly from states in $S_k$. Let $\mathbf{M}_k$ be a matrix such that $(M_k)_{ts}$ is the rate parameter for the transition from $s \in S_k$ to $t \in T_k$. The basis for computing these sets and matrices is shown in **Supplemental Table S1**.

Let $\mathbf{P}_k$ be the corresponding transition probability matrix:

$$(P_k)_{ts} := \frac{(M_k)_{ts}}{\sum_{t' \in T_k} (\mathbf{M}_k)_{t's}}. \tag{3}$$

Let $\mathbf{L}_k$ be the expected time until the next event for every state in $S_k$:

$$(L_k)_s := \frac{1}{\sum_{t \in T_k} (\mathbf{M}_k)_{ts}}. \tag{4}$$

The expected TMRCA for all $s \in S_k$, $\mathbf{d}_k$, is given by the expected time to transition to the next state, $\mathbf{L}_k$, plus the expected time from the next state to the root, which can be expressed as $\mathbf{P}_k \mathbf{d}_{\downarrow k}$, where $\mathbf{d}_{\downarrow k}$ gives the expected time to the root for all possible states that follow GSBL $k$. Therefore

$$\mathbf{d}_k = \mathbf{L}_k + \mathbf{P}_k \mathbf{d}_{\downarrow k}. \tag{5}$$

We will use the superscript 0 to indicate no change in GSBL (remaining at $k$) and the superscript $-$ to indicate a reduction in GSBL (lowering to $k-1$ or $k-2$). Then the set of next states $T_k$

[1] Note that it is possible to skip $k-1$ and transition directly to $k-2$ in the case of a purple-purple coalescence.

can be broken into $T_k^0 = T_k \cap S_k$ and $T_k^- = T_k \setminus T_k^0$. Likewise $\mathbf{P}_k$ can be broken up as $\mathbf{P}_k^0$ (whose target states have GSBL $k$) and $\mathbf{P}_k^-$ (whose target states have GSBL $< k$). The vector $\mathbf{d}_{\downarrow k}^-$ will correspond to the expected TMRCA starting from the states of GSBL $< k$, while $\mathbf{d}_{\downarrow k}^0$ or the expected times for states with GSBL $k$, is simply $\mathbf{d}_k$. Then equation 5 becomes

$$\mathbf{d}_k = \mathbf{L}_k + \mathbf{P}_k^0 \mathbf{d}_k + \mathbf{P}_k^- \mathbf{d}_{\downarrow k}^-, \tag{6}$$

which is equivalent to

$$\mathbf{d}_k = \left(\mathbf{I} - \mathbf{P}_k^0\right)^{-1} \left(\mathbf{L}_k + \mathbf{P}_k^- \mathbf{d}_{\downarrow k}^-\right). \tag{7}$$

Similarly for TSBL,

$$\mathbf{w}_k = \left(\mathbf{I} - \mathbf{P}_k^0\right)^{-1} \left(k \cdot \mathbf{L}_k + \mathbf{P}_k^- \mathbf{w}_{\downarrow k}^-\right). \tag{8}$$

It is therefore possible to solve for $\mathbf{d}_k$ and $\mathbf{w}_k$ from $\mathbf{d}_{k'}$ and $\mathbf{w}_{k'}$ for $k' < k$ by a dynamic programming method, building from $\mathbf{d}_1 = \mathbf{w}_1 = \mathbf{0}$, and computing the relevant matrices and doing the matrix multiplication for each successive value of $k$.
